# MasterPATH: network analysis of functional genomics screening data

**DOI:** 10.1101/264119

**Authors:** Natalia Rubanova, Anna Polesskaya, Anna Campalans, Guillaume Pinna, Jeremie Kropp, Annick Harel-Bellan, Nadya Morozova

**Affiliations:** Université Paris Diderot, Sorbonne Paris Cité, 75013 Paris, France; Skolkovo Institute of Science and Technology, Skolkovo, Russia; Ecole Polytechnique, Université Paris-Saclay, CNRS UMR 7654, Palaiseau, France; Institute of Molecular and Cellular Radiobiology, CEA, F-92265 Fontenay-aux-Roses, France.; INSERM, U967, F-92265 Fontenay-aux-Roses, France.; Université Paris Diderot, U967, F-92265 Fontenay-aux-Roses, France.; Université Paris Sud, U967, F-92265 Fontenay-aux-Roses, France.; Institute for Integrative Biology of the Cell (I2BC), IBITECS, CEA, CNRS, University Paris XI, University Paris-Saclay, 91198, Gif-sur-Yvette cedex, France; Institut des Hautes Etudes Scientiques, 91440 Bures-sur-Yvette, France

## Abstract

Functional genomics employs several experimental techniques to investigate gene functions. These techniques such as loss-of-function screening and transcriptome profiling performed in a high-throughput manner give as result a list of genes involved in the biological process of interest. There exist several computational methods for analysis and interpretation of the list. The most widespread methods aim at investigation of biological processes significantly represented in the list or at extracting significantly represented subnetworks. Here we present a new exploratory network analysis method that employs the shortest path approach and centrality measure to uncover members of active molecular pathways leading to the studied phenotype based on the results of functional genomics screening data. We present the method and we demonstrate what data can be retrieved by its application to the terminal muscle differentiation miRNA loss-of-function screening and transcriptomic profiling data and to the ‘druggable’ loss-of-function RNAi screening data of the DNA repair process.

## Introduction

The most wide-spread functional genomics techniques are loss-of-function screening and transcriptome profiling. The biological mechanism for repression of gene expression by causing the degradation of mRNA is exploited in the RNA interference(RNAi) based loss-of-function screening. Synthesized sequence specific RNAi reagents are introduced into the cells in either arrayed or pooled experiment(Mohr et al. 2014). Typically, small interfering RNAs (siRNAs) or short hairpin RNAs (shRNA) are used for mammalian cells(Mohr et al. 2014). Within the cell, shRNAs are recognized by the Dicer complex, which cleaves the shRNAs into the siRNAs, while siRNAs are directly incorporated into RISC (RNA induced silencing) complex(Sontheimer 2005). Next, RISC-siRNA complex binds to a complementary nucleotide sequence of a protein-coding mRNA transcript. This interaction allows a nuclease in the RISC complex to cleave and to destroy the protein-coding mRNA, therefore silencing expression of the gene in a relatively sequence-specific manner(Sontheimer 2005).

In contrast to RNAi technique that allows to perform only temporal gene knock down, permanent gene knockout can be archived using CRISPR/Cas9 technique. CRISPR-Cas system is a bacteria and archaea adaptive immunity mechanism(Wiedenheft, Sternberg, and Doudna 2012). In nature Cas9, crRNA and tracrRNA are transcribed from the CRISPR locus and form crRNA:tracrRNA:Cas9 complex, where crRNA:tracRNA hybrid acts as a guide which allows endonuclease Cas9 to cleave in a sequence specific manner foreign genetic material(Jinek et al. 2012). A synthesized CRISPR library consists of chimeric RNA produced by fusing crRNA and tracrRNA(Mali et al. 2013; Cho et al. 2013). These RNAs known as sgRNAs form sgRNA:Cas9 endonuclease complex that is transfected into the cell(Mali et al. 2013). Whatever technique is used for silencing the gene, the result of the silencing is assessed with a reporter assay specific to the studied phenotype(Mohr et al. 2014). The development of siRNA(Sui et al. 2002; Sen et al. 2004; Yu, DeRuiter, and Turner 2002; Chung et al. 2008; Yang et al. 2006), shRNA(Hu and Luo 2012; Yu, DeRuiter, and Turner 2002), CRISPR sgRNA(Shalem et al. 2015; Tim Wang, Lander, and Sabatini 2016; T Wang et al. 2014) libraries targeting whole genomes made it possible to perform genome scale loss-of function screening.

Transcriptome profiling allows to study a cellular system at the transcriptome level and aims at monitoring gene expression. The most widespread techniques for transcriptome profiling are microarrays and RNA-seq techniques (Lowe et al. 2017). mRNA abundance is measured by hybridization of fluorescently labelled transcripts to short DNA fragments, known as ″probes″, in microarray-based experiment (Lockhart et al. 2000). The probes are arrayed on a solid substrate that is usually made from glass(Lockhart et al. 2000). RNA-seq refers to the combination of a high-throughput sequencing methodology with computational methods to measure the presence of transcripts in an RNA extract(Lowe et al. 2017).

Numerous computational methods to interpret functional genomics datasets and infer molecular machinery underlying a given biological process have been developed in the past 15 years. The most widespread methods can be grouped into two categories. The first category is the pathways analysis methods which can be considered as the gold standard approach for analysis of both transcriptome profiling and loss-of-function screening results. The classical representatives of the pathways analysis methods are over representation analysis (ORA) methods. An ORA method uses a statistical test to assess the enrichment of a list of genes in annotated gene sets or canonical pathways. ORA evaluates the number of the hit genes that belong to a pathway. Then, each pathway is assigned a p-value to show how significantly it is represented in the hit list by testing over or under-representation. The most commonly used statistical tests are based on the hypergeometric, Fisher′s exact, chi-square, or binomial tests (Khatri et al. 2012). Several improvements over the standard ORA were developed including functional class scoring approaches that aims at detecting coordinated changes in pathways (Khatri et al. 2012) and topology-based approaches that consider pathway topology, connectivity and interactome information(Bankhead et al. 2009; Draghici et al. 2007).

The second category is network analysis methods which use molecular interaction networks as a complementary information(Markowetz 2010). Such methods can help to find functionally related biological components in a functional genomics dataset. This can be done by introducing network-based scoring methods that can use ″guilt by association″ principle and information from both network topology and screening results(L. Wang, Tu, and Sun 2009); by introducing the use of the connectivity of subgraphs of protein-protein interaction networks(Kaplow et al. 2009); by using network neighbor information(Jiang et al. 2015); by performing networks functional analysis that relies on assessing the clustering of selected nodes on the network(Cornish and Markowetz 2014); by extracting the largest connected component of a subnetwork that is created from the optimal number of the top-ranked genes(Kairov et al. 2012).

Another way to use molecular interaction networks is to find significantly enriched subnetworks within a functional genomics dataset. Even manual investigation of such subnetworks can give biologically meaningful results(Krishnan et al. 2011; Warner et al. 2014). The molecular interactions networks can be integrated with other types of biological information to archive higher network specificity: with canonical pathways(Tu et al. 2009; De Maeyer et al. 2015); with regulatory interactions(Lan et al. 2011; De Maeyer et al. 2015); with phosphorylation(Nizard et al. 2014). Moreover, subnetworks can be analyzed for finding functional modules(Dittrich et al. 2008; Mitra et al. 2013).

Here we present a new network analysis method to analyze functional genomics datasets. The method aims at elucidating members of molecular pathways leading to the studied phenotype using the results of the functional screening data. The method works on a network that represent human interactome. The network is constructed from 8 databases: HIPPIE (Schaefer et al. 2012), Signor (Surdo et al. 2017), SignaLink (Fazekas et al. 2013), tFacts (Essaghir et al. 2010), KEGG Metabolic Pathways (Ogata et al. 1999), transMir (J. Wang et al. 2009), mirTarBase (Hsu et al. 2011). The method extracts subnetwork built from the shortest paths between hit genes to so called ″final implementers″ - genes that are involved in molecular events responsible for final phenotypical realization (if known) or between hit genes (if ″final implementers″ are not known). The method calculates centrality score for each node and each linear path in the subnetwork as the number of paths found in the previous step that pass through the node and the linear path. Then the statistical significance of each centrality score is assessed by comparing it with centrality scores in subnetworks built from the shortest paths for randomly generated hit lists preserving the degree distribution of the initial hit list. We hypothesis that the nodes and the linear paths with statistically significant centrality score can be considered as putative members of active molecular pathways leading to the studied phenotype.

We illustrate the application of the method by analyzing the results of transcriptomic profiling and miRNA loss-of-function screening of terminal muscle differentiation and of ′druggable′ loss-of-function screening of DNA repair processes.

## Materials and Methods

### Databases

The following databases were used to construct human integrated interactome of molecular interactions:

- Human Integrated Protein-Protein Interaction rEference (HIPPIE),
- Signor,
- SignaLmk,
- TFacts,
- KEGG Metabolic Pathways,
- TransMir,
- MirTarBase.

All databases contain experimentally validated interactions, except SignaLink database which contains a small number of predicted miRNA-mRNA interactions. We used only high confidence interactions from HIPPIE database. The confidence threshold was chosen according HIPPIE documentation(″HIPPIE Howto,″ n.d.). Since all the databases use different types of gene ID, we converted the ids to the HUGO gene nomenclature and we used this nomenclature to construct human integrated network. Singnor database contains some interaction that involve phenotypes, protein families and stimuli, however we used only interactions between proteins, complexes and small molecules. The basic information about the databases is summarized in Table 1.

**Table 1.** Number of nodes, interactions and types of interactions in databases used to construct human integrated network. PPI stands for Protein-Protein Interactions, TF stands for Transcription Factor.

|  | Nodes | Interactions | Types of nodes | Types of interactions | Direction of interactions |
| --- | --- | --- | --- | --- | --- |
| HIPPIE (high confidence) | 9368 | 41520 | proteins | PPI | indirect |
| Signor | 3977 | 13129 | proteins, complexes, small molecules | PPI, enzymatic | direct, indirect |
| Signalink | 3285 | 27295 | proteins, genes, miRNAs | PPI, miRNA-mRNA, TF-gene | direct, indirect |
| tFacts | 2203 | 4312 | TFs, genes | TF-gene | direct |
| KEGG metabolic pathways | 2921 | 8231 | proteins, small molecules | Enzymatic reactions | direct |
| transMir | 324 | 647 | TFs, miRNAs | TF-miRNA | direct |
| mirTarBase | 2269 | 3511 | miRNAs, genes | miRNA-mRNA | direct |

**Table 2.** shows basic topological features of the direct integrated interactome constructed and indirect protein-protein interactome constructed only from. The exponent of the fitted power law distribution in the degree distribution was calculated with powerlaw(Alstott, Bullmore, and Plenz 2014) Python package.

|  | Integrated network | PPI network |
| --- | --- | --- |
| Nodes | 13 419 | 9 365 |
| Interactions | 85 277 | 41 329 |
| Average clustering coefficient | 0.11 | 0.31 |
| Number of connected components | 126 | 121 |
| Average number of neighbors | 9.2 | 8.6 |
| Density | 0.0007 | 0.0009 |
| Diameter | 25 | 12 |
| Average length of the shortest path | 4.5 | 4.1 |
| Exponent of fitted power-law distribution (total degree) | 3.16 | 2.74 |
| Exponent of fitted power-law distribution (in degree) | 3.93 | - |
| Exponent of fitted power-law distribution (out degree) | 2.36 | - |

### MasterPATH algorithm

The following notions are used in the mechanistic model of pathway construction: an unweighted graph *G=(V, E)* represents a network of molecular interactions, where V are nodes that can be proteins, genes, small molecules or miRNAs; E are edges represent molecular interaction between nodes, interactions can be direct or undirect. List of hit genes of size n is as a set *H = {h_i_: h_i_ ∈ V for ∀i ∈[l..n]}.* List of ″final implementers″ of size m as a set *F = {f_i_: f_i_ ∈ V for ∀ | ∈[l..m]}.* A simple linear paths p between a pair of nodes *(v,u): v,u ∈V* is as set of pairs of nodes that represent existing edge in the graph *G: p(v,u)= (v,v_1_),(v_1_,v_2_)*… *(v_n_,u)* where *v_i_ ∈V for ∀i* and each node vi is distinct. Length *L* of the path *p(v,u)* is the number of edges in the path *p.* We distinguish 4 different types of paths:

- Protein-protein paths if all edges represent protein-protein interactions;
- Transcriptional paths if there exist at least one edge that represent transcriptional interaction;
- MiRNA paths if there exist at least one edge that represent miRNA-mRNA interaction;
- Metabolic paths if there exist at least one edge that represent enzymatic reaction.

The algorithm of the method is the following. For a given network *G,* hit list *H,* list of ″final implementers″ Fthe method finds for each pair of hit gene and ″final implementer″ *(h_i_,f_j_)* all the shortest paths *{p_i_}* of four abovementioned types of length less or equal the maximum length *L_max_* (defined by the user) in the network *G.* The search is done using breadth-first algorithm. Then the centrality score which resembles centrality score *c* is calculated for each node *v* and each path *q* (of length of several interactions) as the number of the shortest paths from *{pi}* that pass through the node *v* and the path *q*: c(v)= |p ∈ *{p_i_}*: v ∈ p | c(q) = | p ∈ *{ p_i_}*: q∈ p |. After that, the statistical significance of each score is assessed. 10000 random hit lists are sampled from the set of nodes *N* preserving or not preserving the degree distribution of the initial hit lit. The probability (p-value^Net^) of getting a node *v* or a path *q* with specific centrality score by chance is calculated as a proportion of sampled hit lists for which a node or a short path has the same or greater centrality score.

## Application

### Human muscle differentiation process transcriptomic profiling and functional screening

The screening data from the study by A. Polesskaya et al.(Polesskaya et al. 2013) was taken as the hit list for terminal human muscle differentiation process. In this study, genome-wide miRNA loss-of-function screening on late differentiating human muscle precursor cell line (LHCN) was performed in two-step approach. The primary screening was done in duplicate with LNA antisense inhibitors library targeting 870 miRNAs and a readout assay that detects multinucleated Myosin Heavy Chain (MHC) positive myotubes. Those miRNAs whose depletion resulted in differences to the negative control ≥ 2 SD were selected for the secondary screen which was done in triplicate. The total number of nuclei was checked in addition to the readout assay from the primary screen. A total of 63 miRNAs whose depletion resulted in differences to the negative control ≥ 2 SD were confirmed in the secondary screen.

The screening data from the study by J. Kropp et al.(Kropp et al. 2015) was taken as the second hit list for terminal human muscle differentiation process. Transcriptomic profiling for proliferation and late differentiation stages in human muscle precursor cell line (LHCN) was performed using Affymetrix Human Gene 1.1 ST arrays(Kropp et al. 2015). A total of 571 differentially expressed genes during late differentiation compared to proliferation stage were found with 2-fold change threshold.

Proteins responsible for *inhibition, activation,* and *facilitating* of fusion of myotubes(Lee 2004; Alzhanov, Mclnerney, and Rotwein 2010; Gunning et al. 2001; Y. Wang et al. 2007; Bourmoum, Charles, and Claing 2016; Tachibana and Hemler 1999; Vasyutina et al. 2009; Doherty et al. 2008), namely MSTN, IGF2, ACTA1, MYH1, MYLPF, ARF6, CD81, CD9, CDC42, EHD2, MYOF, were used as a list of ″final implementers″. The analysis was performed in the integrated human interactome.

It was found 2609 the shortest paths of 4 types of length from 2 to 5 interactions from each miRNA in the hit list from loss-of-function screening to each protein in the list of ″final implementers″. The subnetwork constructed from these paths consists of 1063 nodes (384 of which are genes) and 2710 edges without duplicated edges. The centrality score and the p-value were calculated for each node and path in the subnetwork according the procedure described in the *Method* section. 294 paths of length of 3 to 4 interactions, with centrality score ≥ 3 and p-value < 0.05; 240 nodes with centrality score ≥ 3 and with p-value < 0.05 were taken for further analysis. Analysis of the paths with high centrality scores had highlighted a possible role for a number of nuclear receptors (AR, ARDB1, NR3C1) in skeletal muscle differentiation, as well as suggested functions in myogenesis for such proteins as arrestin (ARRB1 and 2), intersectin (ITSN1), the Rho GTP echange factor VAV3, and the teratocarcinoma-derived growth factor (TDGF1). Interestingly, while the IGF1 regulatory role in myogenesis is very well studied, our approach allowed us to include the arrestin proteins in these pathways, and thus elaborate the known IGF1 network in skeletal muscle differentiation. The MEF2D, p300, CCND1 functions in differentiation have been abundantly demonstrated, and their presence among the results serves as a proof of efficiency of the analysis.

It was found 47714 the shortest paths of 4 types and of length from 1 to 5 interactions from each gene in the hit list from transcriptome profiling to each protein in the list of ″final implementers″. The subnetwork constructed from these paths consists of 2847 nodes and 13032 edges without duplicated edges. The centrality score and the p-value were calculated for each node and each path in the subnetwork. 1623 paths of length of 3 to 4 interactions and centrality score ≥ 3, with p-value < 0.05 and 328 nodes with centrality score ≥ 3 and p-value < 0.05 were taken for further analysis. There are 18 miRNAs among these 328 nodes. 4 of them (hsa-mir-125b, has-mir-133a, has-mir-133b, has-mir-145) are the hit miRNAs in the loss-of-function screen. 5 with the highest centrality score are hsa-mir-125b, hsa-mir-29a, hsa-mir-371, hsa-mir-216a, hsa-mir-1. These miRNAs except hsa-mir-371 were shown to be involved in muscle differentiation and/or proliferation(Callis, Chen, and Wang 2007; Winbanks et al. 2011; Meyer et al. 2015; Crist and Buckingham 2009). Moreover, almost half of 18 miRNAs are known to participate in terminal muscle differentiation, and potential roles in myogenesis could be predicted for other miRNAs in this list e.g that regulate cellular proliferation (such as miR-132, miR-145 or miR-224), as well as cardiac hypertrophy (miR-378). Interestingly, the majority of these miRNAs were not found in the original loss-of-function screen, most likely due to the redundancy of miRNA family members. Indeed, as the miRNAs of the same family share the seed sequence, an efficient loss-of-function screen should have contained not only individual miRNA inhibitors, but also the inactivators of whole miRNA families, in order to avoid false negative results. In this sense, our analysis of these data has been important in supplementing a group of miRNA targets that could have been overlooked. This possibility is highlighted by the presence of known myogenesis regulatory miRNAs (miR-1, miR-29a, miR-216a) in the list resulting from the analysis, whereas they have not been picked up by the original experimental screen.

The analysis of paths allowed to identify a potentially novel and interesting pathway in regulation of myogenesis, involving the Myc-associated factor X (MAX), the clathrin-coated pathway regulatory protein AP2M1, and the EH-domain protein EHD2, which links the clathrin coated transport to actin cytoskeleton, and also binds to myoferlin, a factor promoting myotube fusion. Together with integrin subunits ITGA4 and ITGB1, the extracellular matrix component fibronectin (FN1), and the protein chaperon HSP90, these proteins indicate a possible involvement of specific protein transport pathways in terminal myogenic differentiation. In addition, there is a possibility of involvement of beta-catenin (CTNNB1), C-KIT and PRKC in these processes. It should be noted that these three regulatory factors, while extensively studied in a multitude of biological models, have never been shown to be specifically implicated in skeletal myogenesis. Two major regulatory molecules of skeletal myogenesis, MYOD and SMAD3, have been highlighted, together with their previously known muscle-related targets (TGFB1, CDC42, CTCF). Also, they are linked to a number of proteins that have not been previously studied in the context of muscle differentiation (MAX, KIT, AP2M1…).

Next, we compared the lists of nodes and the lists of paths from two experiments. We found 29 nodes and 16 paths common for both loss-of-function screening and transcriptomic profiling. Among the nodes with the highest centrality score, three – IGF1R, CBL, E2F1 – have been suggested to play key roles in the growth, development, and differentiation of skeletal muscle(Fernández et al. 2002; Nakao et al. 2009; Zappia and Frolov 2016). The path with the highest score consists of CDKN1A, MDM2, TCAP, MSTN proteins. The interaction between MDM2 and TCAP is known to be important for cardiac hypertrophy(Tian et al. 2006), it was also shown that TCAP controls secretion of MSTN(Nicholas et al. 2002). Our analysis shows that this path might be activated by the depletion of hsa-mir-17, hsa-mir-106a, hsa-mir-125a, hsa-mir 145, hsa-mir-93. I can also be noted, that not only androgen receptor (AR), but also the estrogen receptor ESR1 can play a role in human skeletal myogenesis. Interestingly, specific integrins (ITGB1, ITGA6) and adaptor proteins (CRKL) have also been found, confirming the importance of certain membrane/adherence structures. Strikingly, both the receptor of activated C kinase (RACK1), and the inhibitor of this kinase (YWHAB, a 14-3-3 protein), as well as multiple other protein-processing enzymes (peptidase inhibitor SERPIN1, casein kinase CSNK1A1, proprotein convertase FURIN, and activator or protein secretion CHRM3) were found by the analysis, attracting the attention to the role of protein metabolism in myogenesis. It was also very interesting to see the chromosome breakpoint generation factor FRA11B among these potential novel factors that might impact on the differentiation of human myoblasts. This comparison has shown potentially novel paths originating from well-known actors in muscle differentiation (such as IGF1R – RACK1 – CD81); and vice versa, has shown that previously unknown potential regulators of myogenesis, such as YWHAB or FRA11B, can act upon proteins that are well known to regulate myotube hypertrophy and/or fusion (IGFR1, CD81).

The fact that the comparison resulted only in a few number of paths might indicate, that although these experimental systems study one biological process, they interfere biological machinery on two different levels – on the level of translation (miRNAs) and on the level on transcription (transcriptome), which might account on two different regulation mechanisms.

We also found that 14 paths from the analysis of transcriptomic profiling have miRNAs hits from the loss-of-function screening and 60 paths from the analysis of loss-of-function screening have hit genes from the transcriptomic profiling on them. These are the paths from the analysis of transcriptomic profiling that pass through hsa-mir-125b which controls IGF2 gene and the paths that pass through has-mir-145 that control TRIP10 protein which, according to OMIM database, has highest expression in skeletal muscle(McKusick-Nathans Institute of Genetic Medicine, Johns Hopkins University (Baltimore, n.d.) and interacts with CDC42 protein. Also, analyzing these paths one can notice the factors participating in at least three major cellular pathways, that, however, have not been extensively studied in skeletal muscle differentiation. These factors include beta-transduction repeat containing protein (BTRC), which has a strong impact on both beta-catenin and NF-kappa B signaling, as well as the p53-related protein TP73, and, finally, the protein LRIG1 that has a strong negative effect on the expression of epidermal growth factor receptor. These pathways represent interesting new directions to follow, in order to further understand the mechanics of skeletal myogenesis.

### Human DNA repair process functional screening

We used the data from the siRNA functional screening that identified 'druggable' genes involved in oxidative damaged DNA repair(Guyon et al. 2015; Robinson et al. 2017). The screening was performed on genetically engineered HeLa cells that express OGG1-GFP fusion protein. OGG1 is a DNA glycosylate protein that is recruited to chromatin to initiate the repair of oxidized chromatin(Guyon et al. 2015). Each ′druggable′ gene was targeted by 3 siRNAs(Robinson et al. 2017). The intensity of chromatin-bound OGG1-GFP was detected after inducing DNA damage and 18 hit genes were identified.

All 18 hit genes were used as a list of ″final implementers″, since little is known about the proteins involved in the final steps of the DNA repair process. The analysis was performed in the protein-protein human interactome.

It was found 4876 the shortest protein-protein paths of length from 1 to 4 interactions from each gene product in the hit list to each gene product in the list of ″final implementers″. The subnetwork constructed from these paths consists of 381 nodes and 1764 edges without duplicated edges. The centrality score and the p-value were calculated for each node and path in the subnetwork according the procedure described in the *Method* section. 64 paths of length of 2 to 3 interactions, with centrality score ≥ 3 and p-value < 0.05; 28 nodes with centrality score ≥ 3 and with p-value < 0.05 were taken for further analysis.

The top 10 nodes with the highest centrality score are histones 3H proteins: HIST1H3A, HIST1H3B, HIST1H3C, HIST1H3D, HIST1H3E, HIST1H3F, HIST1H3G, HIST1H3H, HIST1H3I, HIST1H3J. It is known that the DNA damage is associated with higher level of chromatin mobility(Strzyz 2017; Dion et al. 2012; Miné-Hattab and Rothstein 2012) and it was shown recently that the increase in chromatin mobility is governed by the proteasome-mediated degradation of core histones(Hauer et al. 2017). Other proteins with high centrality score are SETDB1 protein – a member of the SETI family of proteins; WDR5 protein – a core component of SETI family complexes(Ruthenburg et al. 2016); TP53BP1 – a binding partner of the tumour suppressor protein p53. SETDB1 and WDR5 are associated with post-translational histone modification which allows recruitment of the chromatin-associated proteins and protein complexes(Schultz et al. 2002; Odho, Southall, and Wilson 2010). TP53BP1 protein is known to be an important regulator of the cellular response to DNA double-strand breaks(Panier and Boulton 2013).

Figure 1 presents subnetworks visualized with Cytoscape software(Shannon et al. 2003) for the paths of length 3 interactions with centrality score 3. Figure 1a shows that the method identified that two cohesin proteins SMC3 and SMC1A can act by interacting with RAD21 protein, known cohesin-RAD21 complex(Losada 2014) enriched at DNA double-strand break sites and facilitates recombinational DNA repair(Watrin and Peters 2009). Figure 1b shows possible mechanism of involvement of PSMA1, PSMA3, PSMA4 proteins, all members of the 20S proteasome(Coux, Tanaka, and Goldberg 1996), through interaction with AURKB, Aurora Kinase B(Shu et al. 2003), which in turn interacts with histones H3(Crosio et al. 2002). The path ends with histones H3 – SETDB1 interaction. SETDB1 is a histone methyltransferase that specifically methylates histone H3(Schultz et al. 2002) and is also a member of the hit list. The red arrow shows the direction of this interaction on Figure 1a. Considering this, histones H3 are the proteins where the signal converges from different members of the hit list and we hypothesize that histone H3 can be a ″final implementer″ for this system.

**Figure 1.**
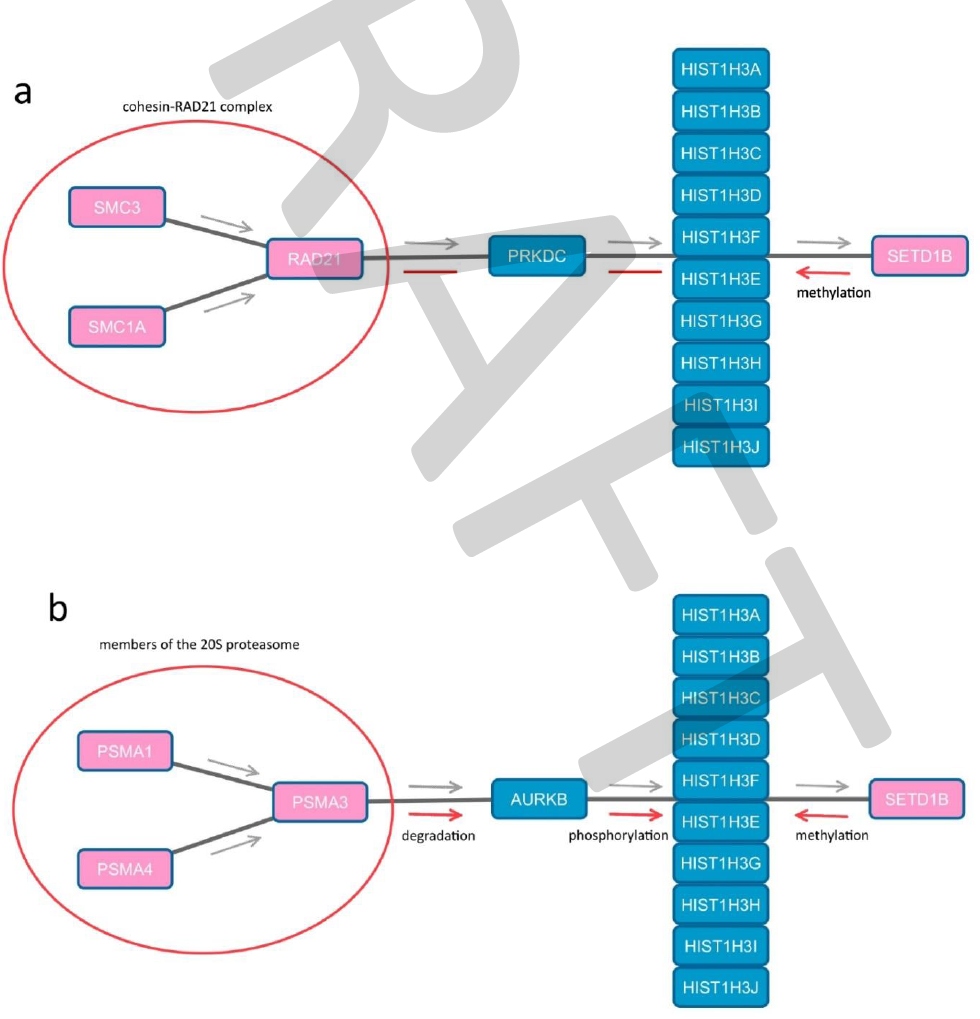
Subnetworks for two paths with centrality score 3. The hit proteins are colored pink, the intermediate proteins are colored blue. The grey arrows show the direction of the paths when it is built by the method. The red arrows show the direction and the effect of the interaction found in literature, **a** RAD21-PRKDC-histone H3-SETDB1 path **b** PSMA3-AURKB-histone H3-SETDB1 path

### Java implementation

The method is implemented in the Java code. The code and full tables for the results presented in the paper available at GitHub page https://github.com/daggoo/masterPath.

## Discussion

We used two different types of networks in our work. The first one was a mixed direct and indirect network constructed from PPI, transcriptional, post transcriptional and metabolic data. The second network was indirect PPI network. PPI networks are the most common networks used in network analysis, although they are known to be incomplete and biased towards the well-studied proteins. Incorporating transcriptional, post-transcriptional and metabolic data, does not solve the issues associated with PPI networks, but adds information on directionality and positive or negative effect of the interactions and gives the ability to build heterogeneous paths.

The bias in the results towards highly connected nodes or paths that pass by highly connected nodes is removed by generating random hit lists. We used the threshold of 0.05 for p-value. However, the nodes/paths with higher p-value can also be considered, it is just not guaranteed that the value of the centrality score is due to the specificity of the node/path to the phenotype or due to its high connectivity.

Other general problems in network analysis is network specificity for the biological system and lack of information relating to some members of the hit list. We used networks that represent human interactome in our work. However, the nodes that are not active can be excluded from the network for some biological systems, based for example on the transcriptomic data, to create a smaller but more specific network. On the other hand, the hit nodes from the functional screening might be poorly studied or might even not be present in the network. Low confidence or predicted interactions for hit nodes might be added to the network in this case.

We introduced a notion of ″final implementer″ in this work. We denoted a ″final implementer″ as a molecule that is involved in events responsible for final phenotypical realization of the biological process. Modern molecular biology accumulated vast amount of knowledge. A part of such molecules is known for some processes, e.g. for apoptosis caspase 3, caspase 6 and caspase 7 could be considered as ″final implementers″. In case such molecules are unknown, the members of the hit list can be used as a list of ″final implementers″ in the analysis on the PPI network and by studying directionality of the paths candidates for ″final implementers″ could be found as we demonstrated for the DNA repair process.

## Conclusion

We presented a new exploratory network analysis method that employs the shortest path approach and centrality measure for uncovering members of active molecular pathways leading to the studied phenotype based on the results of functional screening and in this work. The source code was provided on the GitHub page. We illustrated the application of the method to the analysis of the results of transcriptomic profiling and miRNA loss-of-function screening of human muscle proliferation process and of ′druggable′ loss-of-function screening of human DNA repair process. We also presented the basic topological features of the integrated and the protein-protein human interactome.

